# Tension-dependent stretching and folding of ZO-1 controls the localization of its interactors

**DOI:** 10.1101/156513

**Authors:** Domenica Spadaro, Shimin Le, Thierry Laroche, Isabelle Mean, Lionel Jond, Jie Yan, Sandra Citi

## Abstract

Tensile forces regulate epithelial homeostasis, but the molecular mechanisms behind this regulation are poorly understood. Using structured illumination microscopy and proximity ligation assays we show that the tight junction protein ZO-1 undergoes actomyosin tension-dependent stretching and folding in vivo. Magnetic tweezers experiments using purified ZO-1 indicate that pN-scale tensions (~2-4 pN) are sufficient to maintain the stretched conformation of ZO-1, while keeping its structured domains intact. Actomyosin tension and substrate stiffness regulate the localization and expression of the transcription factor DbpA and the tight junction membrane protein occludin in a ZO-1/ZO-2-dependent manner, resulting in modulation of gene expression, cell proliferation, barrier function and cyst morphogenesis. Interactions between the N-terminal (ZPSG) and C-terminal domains of ZO-1 prevent binding of DbpA to the ZPSG, and folding is antagonized by heterodimerization with ZO-2. We propose that tensile forces regulate epithelial homeostasis by activating ZO proteins through stretching, to modulate their protein interactions and downstream signaling.

---

Both extrinsic mechanical forces, acting on cadherins and integrins, and intrinsic forces, generated by the actomyosin cytoskeleton, control epithelial homeostasis, through the regulation of adhesion and barrier functions, cytoskeletal organization, cell proliferation and morphogenesis ^1–6^. Mechanosensing proteins at adherens and cell-substrate junctions respond to tension by changing their conformation, leading to their enhanced association with actin filaments and actin-binding proteins, and mechanical strengthening of adhesion ^7–10^. Although the polymerization and contractility of the cytoplasmic actomyosin cytoskeleton modulates the nuclear shuttling of transcription factors ^11–14^, it is not clear whether the circumferential actomyosin cytoskeleton associated with tight junctions can regulate signaling by transcription factors. Furthermore, although actomyosin contractility modulates TJ barrier function ^15–17^, it is not known whether any TJ protein can respond to tension by stretching and folding, and whether and how this could affect barrier function.

ZO-1 and ZO-2 ^18, 19^ are cytoplasmic components of TJ, and play a key role in anchoring the junctional actomyosin cytoskeleton to TJ membrane proteins, through the direct interaction of their C-terminal regions with actin filaments ^15, 16^. The N-terminal moiety of ZO-1 and ZO-2 comprises three PDZ domains, followed by Src homology-3 (SH3), U5, and guanylate kinase (GUK) domains ^20^. The PDZ1 and PDZ3 domains bind to the integral TJ membrane proteins claudin(s) and JAM, respectively ^20–25^, and the PDZ2 domain promotes heterodimerization between ZO-1 and either ZO-2 or ZO-3, another member of the ZO family of proteins ^15, 26^. The region of ZO-1 comprising PDZ3, SH3, U5 and GUK domains (ZPSG-1) interacts with the transmembrane TJ protein occludin ^15, 23, 27, 28^, the transcription factor DbpA/ZONAB ^29^, and other ligands (reviewed in ^30^). ZO-1 and ZO-2 are critical for the junctional recruitment and stabilization of both occludin and DbpA, which in turn regulate barrier function ^31–33^, and gene expression, mRNA stability, cell proliferation and survival ^34–36^, respectively. Because of the function of ZO-1 and ZO-2 as linkers between TJ membrane proteins and actin filaments, their conformation might be modulated by tension. In this paper, we provide evidence that mechanical tension generated by the actomyosin cytoskeleton regulates the conformation of ZO proteins and their interactions with DbpA and occludin, to modulate gene expression, cell proliferation, barrier function and epithelial morphogenesis.

## RESULTS

### Structured illumination Microscopy and Proximity Ligation Assay demonstrate force-dependent stretching and folding of ZO-1 in vivo

To test the hypothesis that tensile forces generated by the actomyosin cytoskeleton act on ZO-1 to control its conformation, we expressed ZO-1, tagged with myc and HA epitopes at its N-terminal and C-terminal ends (Fig. 1a), in the context of ZO-1-KO mammary epithelial (Eph4) cells, and we used Structured Illumination Microscopy (SIM) and Proximity Ligation Assay (PLA) to examine the spatial relationships between the tags either in control cells, or in cells depleted of ZO-2, with or without treatment with blebbistatin, an inhibitor of myosin activity. Exogenous ZO-1 was correctly targeted to junctions, and was detected in the cytoplasm of overexpressing cells (Supplementary Fig. 1a-d). In cells treated with a control siRNA, the immunofluorescent signals corresponding to the N-and C-terminal ends of ZO-1 were resolved, regardless of the presence or absence of blebbistatin (Fig. 1b-c). Plotting the intensity of the fluorescent signals as a function of the linear distance across the junction (Fig. 1g) showed that the measured shift between the distribution peaks of green and red fluorescent tags was 107±5.5 nm in cells treated with control siRNA, and 104±7.4 nm in cells treated with control siRNA and blebbistatin (Fig. 1h). This indicates that the distance between the tags of exogenous ZO-1 is not affected by tension in cells expressing ZO-2. The fluorescent signals were also resolved in cells depleted of ZO-2, as long as they were not treated with blebbistatin (Fig. 1e), and the measured shift was 90±19 nm (Fig. 1g-h). In contrast, combining depletion of ZO-2 with blebbistatin treatment resulted in red and green signals that were largely overlapped (Fig. 1f), with a measured shift of 7.42 ±8 nm (Fig. 1g-h). This demonstrates that upon depletion of ZO-2 combined with blebbistatin treatment, ZO-1 undergoes a conformational change, whereby the N-terminal and C-terminal ends come within a closer distance, which does not allow resolution of the signals by SIM. This indicates that in the absence of ZO-2 actomyosin tension is required to maintain ZO-1 in a stretched conformation, and that in the absence of tension ZO-1 undergoes intramolecular folding. Imaging of multicolor microspheres (Fig. 1d) showed a technical chromatic shift in the xy plane ranging from 15 ±13 nm at the center of the field to 55±7 nm at the edges of the field, excluding the possibility the shifts observed between myc and HA tag signals were artifacts. Taking this shift into account, our results indicate that when in the stretched conformation, the length of ZO-1 is between 80 and 120 nm.

**Figure 1:**
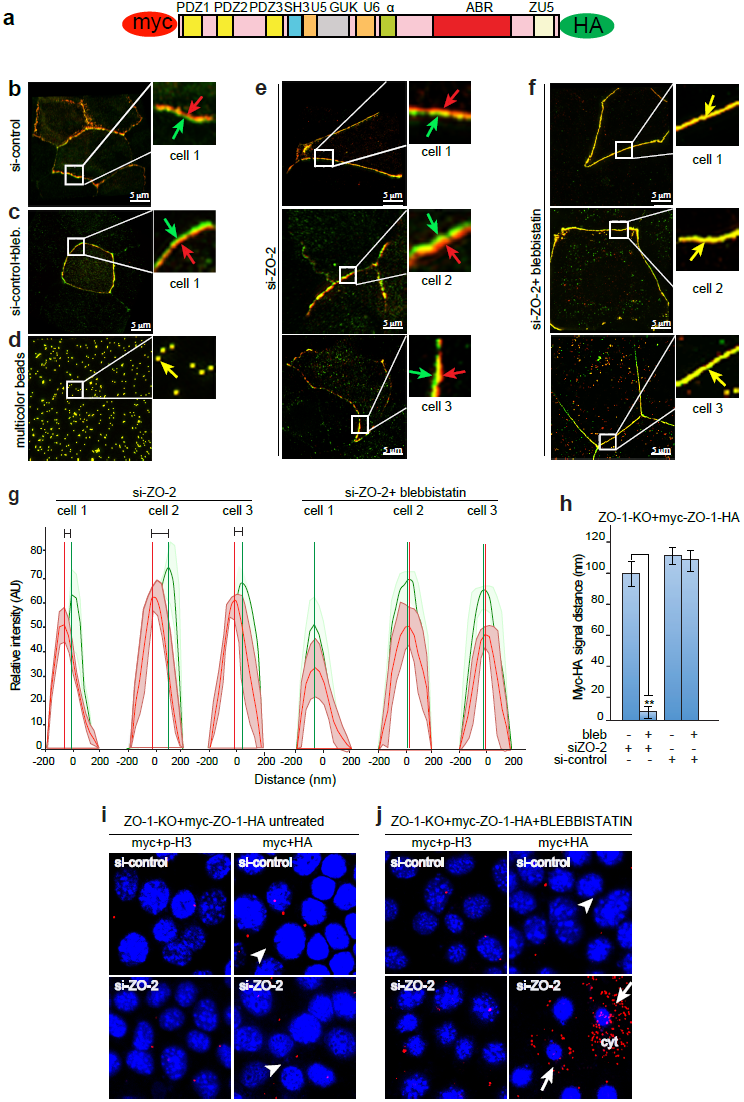
Detection of stretched and folded ZO-1 conformations in vivo by SIM and PLA.

**(a-h) Detection of conformational changes in ZO-1 by SIM**. (**a**) Schematic representation of exogenous myc-ZO-1-HA used to transfect Eph4 cell lines. **(b-c)** SIM images (3D-stacks) of ZO-1-KO Eph4 cells expressing exogenous myc-ZO-1-HA, and treated with control siRNA either in the absence **(b)** or in the presence **(c)** of blebbistatin. **(e-f)** SIM images of ZO-1-KO Eph4 cells expressing exogenous myc-ZO-1-HA, depleted of ZO-2 in the absence of blebbistatin **(e)**, or in the presence of blebbistatin **(f).** Cells were labeled with anti-myc and anti-HA antibodies (red and green signals, respectively), and three representative images for each condition are shown (cell 1, cell 2, cell 3). Green and red arrows in magnified insets **(b, c, e)** indicate spatially resolved labeling of myc (red) and HA (green) epitopes at the N-and C-terminus of ZO-1, respectively. Yellow arrows in **(f)** indicate spatially overlapping signals for myc and HA. **(d)** Multicolor TetraSpeck Fluorescent microspheres (arrow in inset) used as internal control to measure the chromatic shift (xy plane). **(g)** Linescan mean plots (shaded error areas) of fluorescent intensities of fluorophore signals, as a function of distance, corresponding to images of cells in **(e, f)**. The distances (horizontal segments) between vertical lines (maximum intensity peaks) correspond to calculated distances between red and green peaks of fluorescence. **(h)** Histogram plotting calculated distance between peaks of fluorescence intensities of myc and HA epitopes in all SIM images **(b-f** and others not shown), as a function of experimental treatments (indicated below the histogram). Error bars indicate standard error (n=30 linescans for each experimental condition). **(i-j) Detection of conformational changes in ZO-1 by PLA.** Depletion of ZO-2 plus blebbistatin treatment increases proximity between the N-terminal and C-terminal ends of ZO-1. Eph4 ZO-1-KO cells expressing myc-ZO1-HA were either not treated **(i)** or treated with blebbistatin **(j)** and analyzed by PLA. Antibodies for PLA were myc and phospho-histone-H3 (myc + p-H3, negative control), myc and HA (myc+HA, experimental). Top panels=control siRNA. Bottom panels = si-ZO-2. Arrows and arrowheads indicate strong or weak PLA signal, respectively.

To confirm the tension-dependent changes in ZO-1 conformation, we used PLA ^37^ to detect the physical proximity between myc and HA tags of exogeneous ZO-1, under the different experimental conditions (Fig. 1i-j). Little or no PLA signal, comparable to the negative control, was detected between myc and HA in the absence of blebbistatin, regardless of the presence or depletion of ZO-2 (arrowheads in Fig. 1i, myc+HA), indicating that the N-terminal and C-terminal regions of ZO-1 are > 30 nm apart in the absence of blebbistatin. In contrast, in the presence of blebbistatin, the PLA signal between myc and HA tags was dramatically increased upon depletion of ZO-2, and was detectable both at cell-cell junctions, and in the cytoplasm of overexpressing cells (arrows in Fig. 1j, myc+HA, si-ZO-2, and quantification in Supplementary Fig. 1e-f). This indicates that the N-terminal and C-terminal regions of ZO-1 are within 30-40 nm distance when actomyosin contractility and ZO protein heterodimerization are inhibited. In summary, these experiments show conformational changes of ZO-1 in vivo, which indicate that ZO-1 can exist in stretched and folded conformations, depending on actomyosin tension and heterodimerization with ZO-2.

### The structured domains of ZO-1 are mechanically unfolded in vitro at forces > 5pN

To probe the mechanical stability of ZO-1, we purified recombinant, full-length ZO-1 from baculovirus-infected insect cells, and applied force to single molecules, using magnetic tweezers ^38, 39^ (Fig. 2a). When force was linearly increased with a loading rate of 1 pN/s, the domains of ZO-1 were mechanically unfolded at forces of 5-45 pN, as indicated by multiple extension steps (Fig. 2b, red and blue curves). The unfolded domains of ZO-1 could be refolded at forces <5 pN, with a loading rate of -0.3 pN/s (Fig. 2b, black curves, and inset). These results suggest that forces of several pN to tens of pN can cause unfolding of structured domains within full-length ZO-1. Based on the step sizes of all the unfolding events, we estimate that 800-900 residues, which span about 50% of the full-length ZO-1 sequence, and likely correspond to the N-terminal half of ZO-1, are in a stably folded conformation at forces <5 pN. The rest of the molecule, lilely the C-terminal half of the molecule, is either in an intrinsically disordered conformation, or mechanically too weak to be determined in our experiments. Assuming that structured domains occupy half of the residues in full-length ZO-1, and the other half is in a flexible disordered conformation, we estimate that a force of ~ 2-4 pN is needed to maintain the stretched conformation of ZO-1, with an N-to-C distance of ~100 nm, estimated on the basis of SIM experiments (Fig. 1). This force is significantly smaller than that needed to unfold the structured domains (> 5 pN, Fig. 1b).

**Figure 2:**
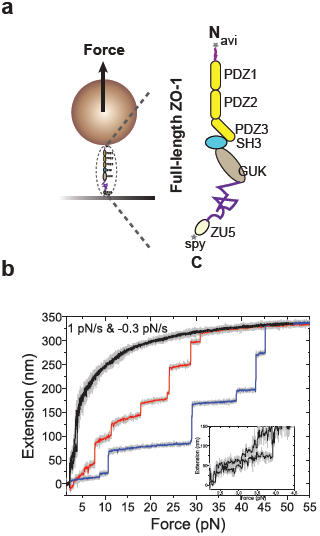
(a-b) Force-dependent unfolding and refolding dynamics of full-length ZO-1.

**(a)** Schematic representation of molecular tweezers experimental configuration (left): ZO-1 molecule was immobilized on the substrate, and force was applied. Schematic domain organization of ZO-1 (right) with structured domains in the N-terminal regions, and disordered C-terminal region. (**b**) Typical force-extension curves of ZO-1 unfolding (red and blue tracings) and refolding (black). Each extension jump of the colored curves indicates an unfolding of a domain or sub-domain during force-increase scans (with a loading rate of 1 pN/s). Bottom Inset shows the zoom-in of refolding events of ZO-1 during force-decrease scans (with a loading rate of -0.3 pN/s). The black or colored lines are 5-points FFT smooth of the raw data (light gray).

### The stretched conformation of ZO proteins promotes the junctional localization of DbpA and occludin in vivo

Next, we explored the biological consequences of ZO-1 stretching and folding. ZO-1 and ZO-2 recruit the transcription factor DbpA and the transmembrane protein occludin to TJ, by binding to them through its ZPSG (PDZ3-SH3-U5-GUK) domain. We asked whether promoting the folded conformation of ZO-1, by depleting cells of ZO-2 and treating them with blebbistatin, impacted on the junctional accumulation of DbpA and occludin (Fig. 3). In the absence of blebbistatin, DbpA was detected at junctions both in WT cells, and in cells depleted of ZO-2 (arrows in Fig. 3a). However, upon treatment with blebbistatin, DbpA labeling was strongly reduced at junctions of ZO-2-depleted cells (arrowheads in Fig. 3b, and quantification in Fig. 3c), but not WT cells (arrows in Fig. 3b, and quantification in Fig. 3c). Immunoblot analysis showed that DbpA levels were reduced in cells depleted of ZO-2 and treated with blebbistatin (red box in Fig. 3d), phenocopying the proteolytic degradation observed in ZO-1-KO cells depleted of ZO-2 ^36^. Similarly, junctional labeling for occludin was normal in WT and ZO-2-KD cells in the absence of blebbistatin (arrows in Fig. 3e), but was dramatically reduced in the presence of blebbistatin, only in ZO-2-KD cells (arrowheads in Fig. 3f). Thus, experimental conditions that lead to the folded conformation of ZO-1 (Fig. 1) result in decreased localization of DbpA and occludin at junctions, indicating that binding between the ZPSG region of ZO-1 and its interactors in inhibited in vivo when ZO-1 is in a folded conformation.

**Figure 3:**
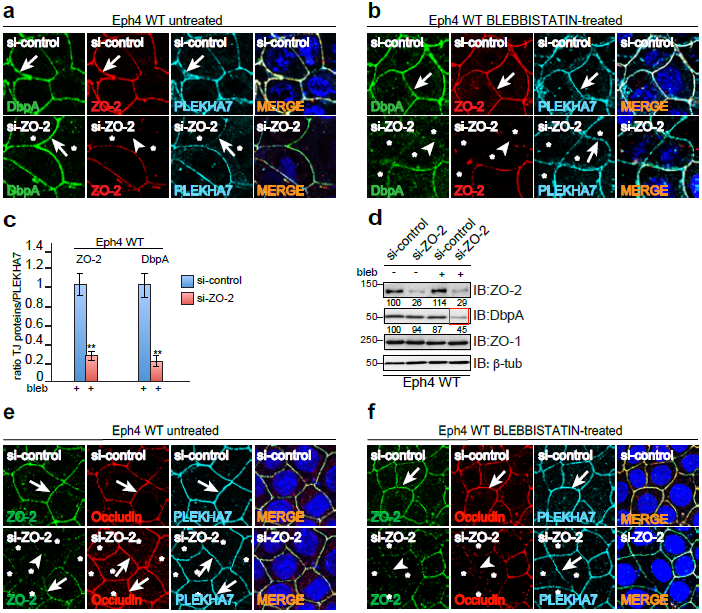
The junctional accumulation of DbpA and occludin, and DbpA stability, are regulated by force in a ZO-1-dependent manner. **(a-c)** Depletion of ZO-2 combined to blebbistatin treatment reduces DbpA accumulation at junctions**. (a, b)** Immunofluorescent localization of DbpA (green), ZO-2 (red, to identify depleted cells), PLEKHA7 (blue, internal reference for junctions) in WT Eph4 cells treated either with si-control (top panels) or si-ZO-2 (bottom panels), either not treated (**a**), or treated with blebbistatin (**b**). Each panel is labeled with the experimental condition (top, si-control or si-ZO-2) and with the antibody used for labeling (bottom), in the color of the fluorophore (green=Alexa488/FITC, red=Cy3, blue=Cy5). Arrows and arrowheads indicate normal or decreased/absent junctional labeling, respectively. Merge panels show nuclei labeled in blue by DAPI. (**c)** Histogram showing a semiquantitative analysis of decrease in junctional labeling 67 for the indicated TJ proteins (ZO-2, DbpA) in WT Eph4 cells treated with blebbistatin, using PLEKHA7 as a reference for junctions. Treatments were with either si-control or si-ZO-2 ((+) or (-) below the histogram), for n=3 separate experiments. Asterisks indicate statistical significance, as determined by Student’s T-test (**, p<0.01). (**d)** DbpA protein levels are reduced in WT cells treated with blebbistatin and depleted of ZO-2. Immunoblot analysis of ZO-2, DbpA, and ZO-1 (and β-tubulin, as a control protein loading normalization) in si-RNA-treated cells (si-control=blue, si-ZO-2=red), either untreated or treated with blebbistatin (bleb, - or +). **(e, f)** Occludin junctional localization is decreased in WT cells depleted of ZO-2 and treated with blebbistatin. Immunofluorescent localization of occludin (red), ZO-2 (green, to identify depleted cells), PLEKHA7 (blue, internal reference for junctions) in WT Eph4 cells treated either with si-control (top panels) or si-ZO-2 (bottom panels), either not treated **(e)**, or treated with blebbistatin **(f)**. Arrows and arrowheads indicate normal or decreased/absent junctional labeling, respectively. Merge panels show nuclei labeled in blue by DAPI.

Since DbpA and occludin interact with both ZO-1 and ZO-2 ^15,23,36,40^, we used ZO-1-KO cells to ask whether tension affects not only ZO-1 dependent, but also ZO-2-dependent accumulation of DbpA and occludin at junctions (Supplementary Fig. 2). Furthermore, in addition to blebbistatin, we tested different drugs that affect the organization of the actin cytoskeleton. Specifically, we treated mixed cultures of wild-type (WT) and ZO-1-KO cells with either latrunculin A, which prevents the polymerization of actin filaments, CK-666, which inhibits Arp2/3-dependent nucleation of branched actin filaments, SMIFH2, which inhibits the formin-dependent nucleation of bundled actin filaments (Supplementary Fig. 2a), and blebbistatin (Supplementary Fig. 2b). DbpA was localized at junctions of ZO-1-KO cells (arrows in Supplementary Fig. 2a-control) when cells were not treated with drugs, consistent with the notion that ZO-2 alone, in the absence of ZO-1, can sequester DbpA at junctions of confluent cells ^36^. However, upon treatment with latrunculin A, SMIFH2, but not CK-666, DbpA was undetectable at junctions of ZO-1-KO, but not of WT cells (arrowheads in Supplementary Fig. 2a – LAT-A, SMIFH2 and arrow in Supplementary Fig. 2a – CK-666). The lack of activity of CK-666 indicated that branched actin filament polymerization is not involved in modulating ZO-mediated junctional sequestration of DbpA. Similarly, treatment of mixed WT/ZO-1-KO cells with blebbistatin induced a loss of junctional DbpA only in cells expressing one ZO protein, but not in WT cells (arrowheads in Supplementary Fig. 2b - BL). Phenocopying what was observed in cells depleted of both ZO proteins ^36^, DbpA protein levels were decreased in ZO-1-KO cells treated with blebbistatin (red box in Supplementary Fig. 2c and quantification in Supplementary Fig. 2d), and normal DbpA levels were rescued by treatment with the proteasome inhibitor MG132 (green box in Supplementary Fig. 2c and quantification in Supplementary Fig. 2d), leading to detection of DbpA in the cytoplasm and nucleus of ZO-1 KO cells (“n” in panel BL/MG, Supplementary Fig. 2b). Occludin was also localized at junctions of both WT and ZO-1-KO cells (arrows in Supplementary Fig. 2e, control), but, upon treatment with blebbistatin, occludin labeling was decreased at junctions of ZO-1-KO, but not WT cells (arrowheads in Supplementary Fig. 2e, BL). Proximity ligation assay (PLA) was used as an additional assay to detect proximity between ZO-1 and occludin (Supplementary Fig. 2f-g). In the absence of blebbistatin, ZO-1 and occludin were in close proximity, regardless of ZO-2 depletion (arrows in Supplementary Fig. 2f). However, in the presence of blebbistatin, ZO-1 was associated with occludin only in the presence, but not in the absence of ZO-2 (arrows and arrowheads in Supplementary Fig. 2g).

In summary, these experiments show that loss of either bundled actin filament organization or myosin activity in cells lacking one ZO protein results in the loss of junctional localization of DbpA and occludin, suggesting loss of interaction with the remaining ZO protein. The results also suggest that, similarly to ZO-1, ZO-2 undergoes tension-dependent stretching, and that stretching activates both ZO-1 and ZO-2, and promotes their binding to DbpA and occludin.

### Tension controls cell proliferation and paracellular barrier function through ZO proteins

To examine the cellular consequences of force-dependent stretching and folding of ZO proteins, we analyzed the signaling outputs by DbpA and occludin under the different experimental conditions.

Translocation of DbpA to the nucleus promotes cell proliferation, enhances transcription of cyclin D1 and PCNA, and represses transcription of ErbB2 ^29,35^. In ZO-1-KO cells treated with blebbistatin and MG132, which show nuclear DbpA labeling (panel BL/MG in Supplementary Fig. 2b) cell proliferation was increased, as determined by labeling with Ki67, and this increase was reduced upon depletion of DbpA (Fig. 4a-b). Under the same conditions, qRT-PCR showed increased expression of the DbpA target genes cyclin D1 and PCNA (Fig. 4c) and decreased expression of ErbB2 (Fig. 4d). These changes in gene expression did not occur in ZO-1-KO cells treated with blebbistatin alone (Fig. 4c-d), conditions under which DbpA is not detected in the nucleus, and DbpA leves are reduced, due to proteasomal degradation (Supplementary Fig. 2b-c).

**Figure 4:**
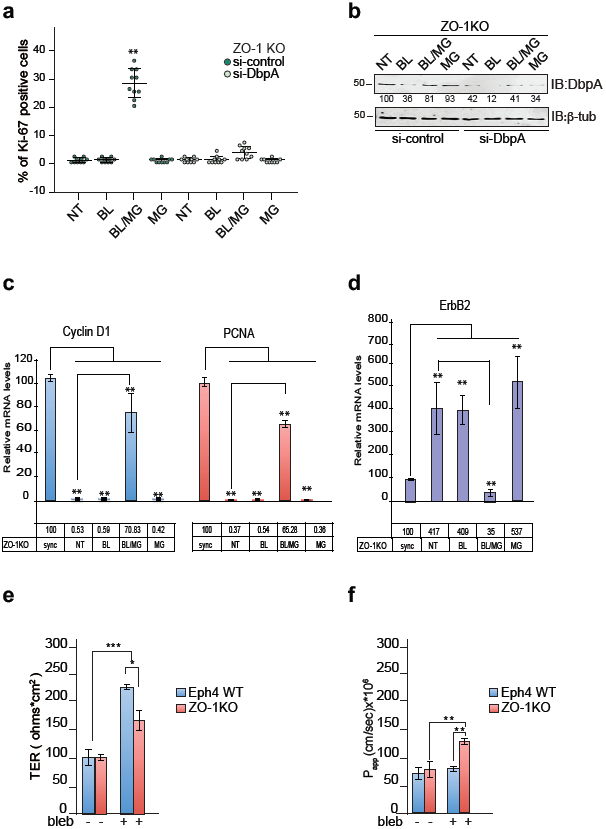
Tension controls cell proliferation and paracellular barrier function through ZO proteins. **(a-b)** The effects of DbpA depletion on cell proliferation depend on tension and ZO proteins. **(a)** Confluent Eph4 ZO-1KO cells were transfected with siRNA-control or siRNA-DbpA and then were treated with the following drugs (NT= not treated, BL=blebbistatin, BL/MG= blebbistatin + MG132, MG=MG132). The dot-plots show the percentage of Ki-67 positive cells. (n=200 cells for each treatment). Note that the percentage of Ki-67 epithelial positive cells was increased upon BL/MG treatment, and this was abolished by DbpA depletion. **(b)** Immunoblotting showing DbpA levels in untreated ZO-1-KO cells or in cells treated with BL, BL/MG and MG. Numbers below the immunoblot indicate the quantification of band intensity by densitometric analysis. **(c-d)** Treatment of ZO-1-KO cells with blebbistatin results in altered expression of DbpA-regulated genes. qRT-PCR analysis of mRNA levels, showing relative expression of Cyclin D1 (blue), PCNA (red) **(c)** and ErbB2 (purple) **(d)** in ZO1-KO cells, following syncronization of proliferating cells in G1 (sync, value taken as 100%) ^36^, and relative values following different experimental treatments (NT= not treated, BL=blebbistatin, BL/MG= blebbistatin + MG132, MG=MG132). Asterisks indicate statistical significance (**, p<0.01), as determined by Student’s T-test (n=3 independent experiments for b-c). WT cells are not shown since DbpA is always junctional. **(e-f)** Tight junction barrier function of ZO-1-KO but not WT confluent monolayers is altered in the presence but not in the absence of blebbistatin. Histograms show transepithelial electric resistance (TER, **(e)**) and permeability to 3 kDa FITC dextran (P, **(f)**) in WT (blue) and ZO-1-KO (red) cells, either untreated (-) or treated with blebbistatin (bleb +). Error bars represent standard deviations (n=3, *= p≤0.05, **= p≤0.01, ***= p≤0.001).

Occludin contributes to regulating TJ barrier function ^31,41,42^. Measurement of TJ barrier function showed that in the absence of blebbistatin no significant differences were observed in permeability either to ions (TER, Fig. 4e) or to larger molecules (P_app_, Fig. 4f), of WT versus ZO-1-KO Eph4 cells (see also ^43^). However, blebbistatin induced an increase in the TER of WT cells, and this increase was significantly lower in ZO-1-KO cells (TER, Fig. 4e). In addition, blebbistatin treatment did not influence the flux of FITC-dextran across monolayers of WT cells, whereas it induced an increased flux in ZO-1-KO cells (Fig. 4f), suggesting that the barrier is more sensitive to actomyosin tension when only ZO-2 is expressed.

Taken together, these results suggest that in cells expressing only one ZO protein the organization and contractility of the actomyosin cytoskeleton, which promotes stretching and activation of ZO proteins, is required to support both DbpA-dependent regulation of gene expression and occludin-dependent modulation of barrier function.

### Substrate stiffness and actomyosin contractility modulate epithelial proliferation and cyst growth in 3D culture through ZO proteins

Since substrate stiffness influences intracellular tension, we examined cyst morphogenesis in 3D cultures of Eph4 cells grown in Matrigel, a softer and more physiological substrate, compared to glass coverslips ^44,45^. WT Eph4 cells formed cysts with a central lumen, and their size increased to an average diameter of 250 μm within 21 days (Fig. 5a,e). In contrast, ZO-1-KO cells formed cysts, which grew slowly in Matrigel, and had an average diameter of 50 ¼m and no lumen after 21 days (Fig. 5c,e). This suggests that under these conditions ZO-2 is folded (inactive), leading to loss of binding and degradation of DbpA, and decreased proliferation and growth. These results also indicate that growth on a soft substrate phenocopies treatment of cells growing on coverslips with blebbistatin. Next, we asked whether this phenotype of ZO-1-KO cysts could be rescued by artificially enhancing actomyosin contractility. To this purpose, we incubated cells in 2’-deoxyadenosine-5-triphosphate (dATP) ^46,47^. Addition of dATP did not affect the growth rate and size of WT cysts (Fig. 5b,e). In contrast, it rescued the stunted growth phenotype of ZO-1-KO cells, resulting in cyst growth and size similar to WT cells (Fig. 5d,e). This suggests that ZO-2 can be activated by stretching, induced by increased actomyosin contractility. Immunoblotting showed that DbpA protein levels were reduced by about 80% in ZO-1-KO cysts, compared to WT, suggesting proteolytic degradation, and this was rescued by treatment with dATP (red and green boxes in Fig. 5f). Immunofluorescence showed that in WT cells DbpA was localized in the nucleus at early time points (Fig. 5g), whereas at later stages it was localized at junctions (Fig. 5h), both in the presence and absence of dATP. In contrast, DbpA was undetectable by immunofluorescence in ZO-1-KO cysts both at early (Fig. 5i) and late (Fig. 5j) time points. Significantly, dATP rescued both the expression (Fig. 5f) and the localization of DbpA at all time points (Fig. 5i-j), correlating with increased cyst growth (Fig. 5d). Labeling for E-cadherin showed expression and localization at cell– cell contacts in both WT and ZO-1-KO cells, independently of dATP, suggesting that cells lacking ZO-1 can still form adherens junctions (Fig. 5g-j).

**Figure 5:**
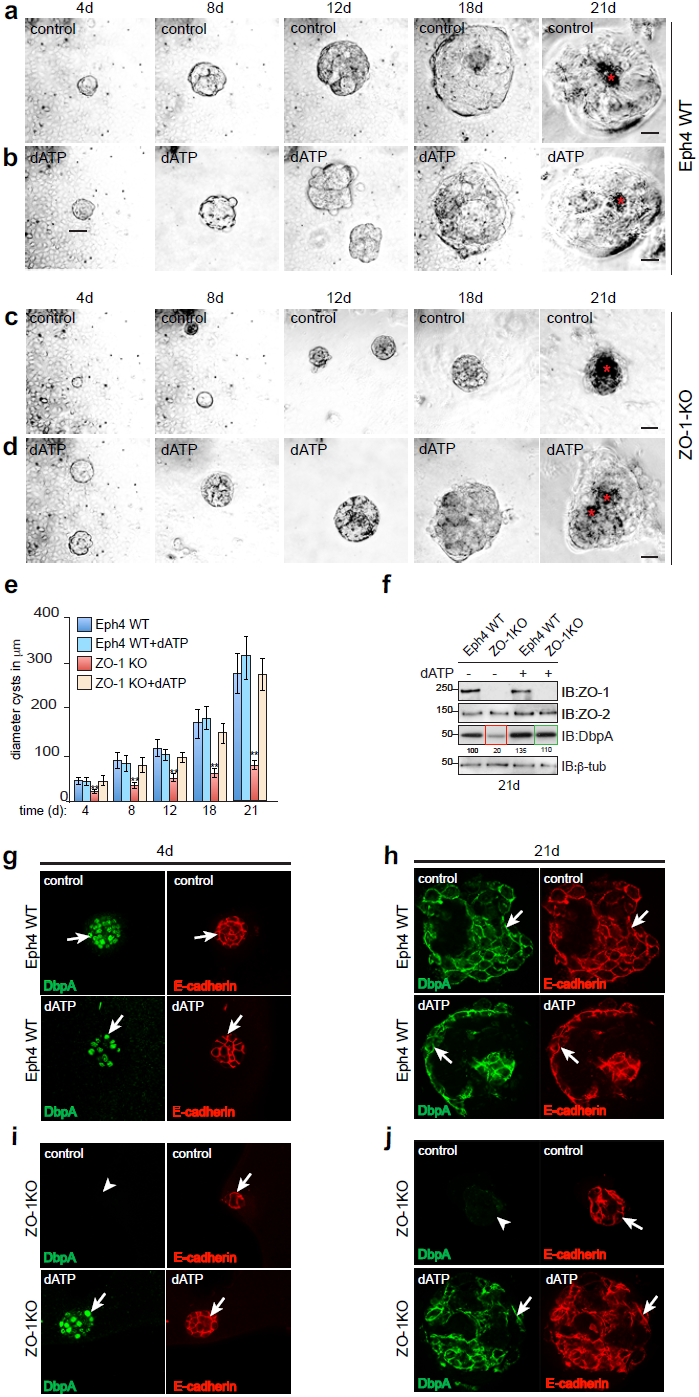
ZO-1 is required for cell proliferation, cyst growth and Dpa accumulation at junctions in Eph4 cells grown in Matrigel. **(a-d)** The growth of ZO-1-KO Eph4 cell cysts is tensio-dependent. **(a and c)** Brightfield microscopic images of cysts from untreated Eph4 WT **(a)** or ZO-1-KO **(c)** cells, at different times (4 days to 21 days) after plating, showing reduced growth of ZO-1-KO cysts. **(b, d)** Cell growth is rescued by dATP treatments in ZO-1 KO cysts. Eph4 WT **(b)** and ZO-1KO **(d)** cysts were treated with dATP after plating. Asterisks indicate lumens. Scale bars =50 μm. **(e-f)** Histograms showing average cyst diameter **(e)** for WT (blue) WT+dATP (light blue), ZO-1-KO (red), and ZO-1-KO (yellow) cysts. Data are from n=30 cysts, from three independent experiments. Error bars represent standard deviations. ** p≤ 0.01. **(f-j)** DbpA levels are decreased in ZO-1-KO cells grown in Matrigel. **(f)** Immunoblot analysis of lysates of either untreated or dATP-treated WT or ZO-1-KO cysts (at 21 days after plating), using antibodies against ZO-1, ZO-2, and DbpA (β-tubulin used for normalization). Numbers on the left in indicate migration of molecular size markers (kDa). Numbers below each lane indicate densitometry quantification of the protein. The red box indicates decreased DbpA protein levels in ZO-1-KO cysts. The green box indicates rescued DbpA protein levels upon treatment of ZO-1-KO cysts with dATP. **(g-j)** Immunofluorescent localization of DbpA and E-cadherin (to label junctions) in untreated or dATP-treated WT or ZO-1-KO cysts, at days 4 **(g, i)**, and 21 **(h, j)** after plating in Matrigel. Arrows and arrowheads indicate normal and reduced/absent staining, respectively. Scale bars=10 μm.

### Intramolecular interactions within ZO proteins prevent their binding to DbpA and occludin, and are inhibited by heterodimerization

Folding of ZO proteins might be the result of an intramolecular interaction between their C-terminal and N-terminal regions, which masks the DbpA and occludin binding sites on the ZPSG domains. To test this hypotheses, we first established which ZO protein interacts with DbpA and occludin in in vitro assays. Either GST-DbpA or GST fused to the C-terminal region of occludin were used as baits, and GFP-tagged ZPSG regions of ZO-1 (ZPSG-1, residues 417-806), ZO-2 (ZPSG-2, 509-880), and ZO-3 (ZPSG-3, 369-750) were used as preys (Supplementary Fig. 3a). DbpA interacted with ZPSG-1 and ZPSG-2, but not with ZPSG-3 (Supplementary Fig. 3b), whereas all ZPSG regions interacted with GST-occludin (Supplementary Fig. 3c).

Next, we focused on ZO-1 and ZO-2, and asked whether intramolecular interactions between their ZPSG and C-terminal regions can occur, using GST pulldown assays (Fig. 6 a-b for ZO-1 and Fig. 6c-d for ZO-2). Different constructs of the C-terminal region of ZO-1 interacted with ZPSG-1, the strongest signal being observed with the most C-terminal fragment of ZO-1 (1619-1748) (Fig. 6b).Similarly, the bacterially expressed C-terminal region of ZO-2 (890-1190) interacted with the ZPSG-2 region (Fig. 6d).

**Figure 6:**
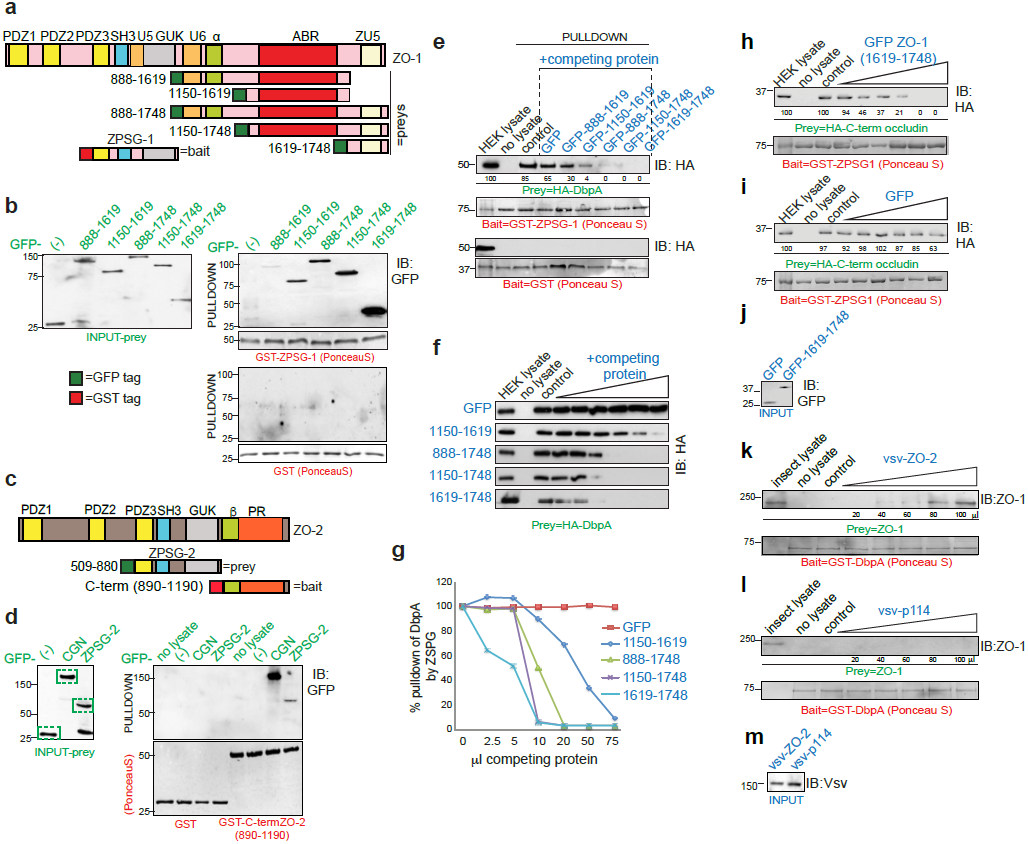
Intramolecular interactions between ZPSG and C-terminal domains of ZO proteins prevent interaction with DbpA. **(a-d)** ZO proteins undergoe intramolecular interactions. **(a)** Schematic diagrams of the domain organization of ZO-1 and of bait and preys used in GST pulldown experiments. **(b)** Immunoblot analysis, using anti-GFP antibodies, of GST pulldowns, using either GST or GST-ZPSG-1 as baits, and GFP-tagged C-terminal fragments of ZO-1 as preys. GFP alone (-) was used as a negative control prey. Normalization of GFP-tagged preys in HEK293T lysates was carried out by immunoblotting with anti-GFP antibodies (INPUT-prey, **(b)**). **(c-d)** ZO-2 undergoes intramolecular interaction.**(c)** Schematic diagrams of the domain organization of ZO-2 and bait and prey used for GST pulldown experiments. **(d)** Immunoblot analysis, using anti-GFP antibodies, of normalized preys (left) and of GST pulldowns, using either GST or GST fused to the C-terminal fragment of ZO-2 (890-1190) as a bait, and the GFP-tagged ZPSG-2 region as a prey. GFP-cingulin (CGN) was used as a positive control. Normalization of GFP-tagged preys in HEK293T lysates was carried out by immunoblotting with anti-GFP antibodies (INPUT-prey, **(d)**). Bait normalization (GST-ZPSG-1 **(b)**, GST-ZO-2(890-1190) **(d)**) was carried out by PonceauS staining of blots. Pulldowns are labeled on top with prey constructs and on bottom with baits. Numbers on the left of immunoblots indicate migration of molecular size markers (kDa). Dotted green boxes in **(d)** show prey proteins (normalized inputs). Data are representative of > n=3 independent experiments. **(e-g)** C-terminal fragments of ZO-1 inhibit ZPSG-1 interaction with DbpA. **(e)** Pulldowns using either GST-ZPSG-1 (top) or GST (bottom, negative control) as baits and HA-DbpA as prey, either in the absence, or in the presence of competing proteins (annotated in blue): either GFP, or GFP fused to C-terminal fragments of ZO-1 (amino acids of each construct indicated above the blot). Interacting HA-DbpA is visualized by immunoblotting with anti-HA antibodies, and is decreased in the presence of competing ZO-1 fragments. **(f)** Immunoblot showing titration of competition pulldowns using increasing amounts of either GFP, or the indicated GFP-tagged fragments of ZO-1 C-terminal region (indicated on the left). **(g)** Plot showing quantification (by densitometry analysis of immunoblots) of HA-DbpA bound to ZPSG-1 from the experiment shown in **(e)**, as a function of volume of HEK293T lysate containing the competing protein. The amounts of competing proteins were normalized by immunoblotting (Fig. 6b). **(h-j)** Occludin and C-terminal regions of ZO-1 compete for binding to the ZPSG-1 domain. **(h, i)** Immunoblot analyses of pulldowns using GST-ZPSG-1 as a bait, HA-tagged C-teminal occludin (residues 406-521) as a prey, and increasing amounts of either GFP-tagged C-terminal ZO-1 fragment (GFP-ZO-1-1619-1748) **(h)** or GFP alone **(i)** as competitors. Normalization of competing proteins for pulldowns is shown in **(j)**. **(k-m)** Heterodimerization promotes ZO-1 interaction with DbpA. **(k-l)** Immunoblot analysis of GST pulldowns using GST-DbpA as a bait, and full-length ZO-1 as a prey, in the presence of increasing amounts (μl of lysate indicated below each lane) of either vsv-tagged ZO-2 **(k),** or vsv-tagged p114-RhoGEF **(l)**. **(m)** shows normalized prey (INPUT) for pulldowns shown in **(k, l)**. Numbers below immunoblots indicate densitometry quantification of band intensity. Ponceau S staining for bait normalization is shown below immunoblots.

Next, we asked whether the interaction of the C-terminal region of ZO-1 with ZPSG-1 inhibits the binding of ZPSG-1 to either DbpA or occludin. To this purpose, we carried out competition GST pulldown assays, using HA-tagged DbpA as a prey, either in the absence, or in the presence of increasing amounts of competing C-terminal fragments of ZO-1 (Fig. 6e-g). The interaction of DbpA with ZPSG-1 was not inhibited by GFP alone, but was inhibited by GFP-tagged fragments comprising different regions of the C-terminal region of ZO-1, the most C-terminal fragment (residues 1619-1748) being the most effective (Fig. 6e). Importantly, both the ZO-1 C-terminal fragment (1619-1748) and DbpA bind to the SH3 domain within the ZPSG-1 (Supplementary Fig. 4a-b), accounting for their reciprocal inhibition of binding. The calculated dissociation equilibrium constants (K_d_s) for the interaction of ZPSG-1 with the ZO-1 C-terminal fragment (1619-1748) and with DbpA were 66 nM and 100 nM, respectively, on the basis of a supernatant depletion assay and (Supplementary Fig. 4g-j). Similarly, DbpA but not CFP inhibited the interaction between the ZPSG region of ZO-2 (ZPSG-2) and the C-terminal region of ZO-2 (890-1190) (Supplementary Fig. 4d-f). Competition pulldown assays were also carried out usingt he C-terminal region of occludin as a prey (Fig. 6h-j). The GFP-tagged C-terminal region of ZO-1 (1619-1748) (Fig. 6h), but not GFP alone (Fig. 6i) inhibited the interaction between ZPSG-1 and the C-terminal region of occludin. Taken together, these data indicate that interactions between the C-terminal and ZPSG domains within ZO proteins can inhibit binding of the ZPSG domains to DbpA and occludin.

Full-length ZO-1 does not interact with GST-DbpA in pulldown assays ^36^. Since ZO-1 and ZO-2 form heterodimers through their PDZ2 domains ^15, 26^, and heterodimerization promotes the junctional localization of DbpA under low tension conditions (Fig. 1, Fig. 3), we asked whether heterodimerization promotes the interaction of full-length ZO-1 with DbpA in vitro. Using GST-DbpA as a bait, we detected ZO-1 in pulldowns when increasing amounts of vsv-tagged full-length ZO-2 (Fig. 6k), but not vsv-tagged p114-RhoGEF (negative control, Fig. 6l), were added. This suggested that heterodimerization stabilizes the stretched conformation, and makes the ZPSG region available for binding to its ligands in vitro.

## DISCUSSION

The organization of the actomyosin cytoskeleton at cell-cell junctions is critical for barrier function, adhesion, polarity, and differentiation. Mechanical tension affects cell behaviour, as shown by studies revealing key roles for α-catenin, vinculin and myosin in mechanotransduction. In this paper we investigate ZO-1 and ZO-2, which are good candidates for being transducers of junctional tension, since they directly link actin filaments to transmembrane TJ proteins, and are pivotal for junctional actomyosin organization, through their interaction with actin-binding proteins, such as cortactin, cingulin and α-catenin, and with regulators of Rho GTPases, such as Toca-1, ArhGEF11, and others ^15,48–54^.

Our results provide the first evidence that ZO-1 can exist in vivo in different tension-dependent conformations, and that ZO proteins can transduce force into altered binding to physiological interactors, correlating with different effects on cell behaviour. We conclude that in WT cells ZO protein heterodimerization stabilizes the stretched conformation, allowing interaction of the ZPSG regions with DbpA and occludin both at higher (Fig. 7a) and lower (Fig. 7b) levels of actomyosin-dependent tension. In contrast, in cells expressing only one ZO protein, junctional tension is necessary to promote stretching and ZPSG interaction with DbpA and occludin (Fig. 7c). Reducing actomyosin contractility in these cells, either by culture on a soft substrate, or by treatment with actomyosin-active drugs, leads to intramolecular folding of the ZO protein, destabilization of junctional occludin, and proteasomal degradation of DbpA (Fig. 7d). If DbpA degradation is prevented by MG132, DbpA can translocate into the nucleus, to regulate gene expression and promote proliferation (Fig. 7d).

**Figure 7:**
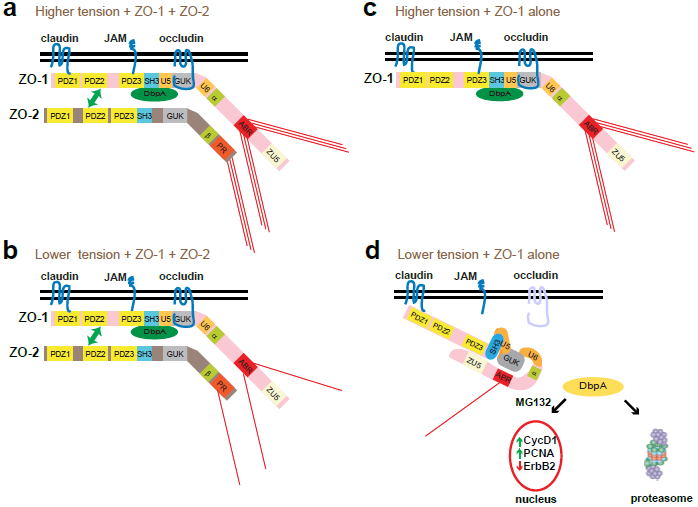
A model for force-and heterodimerization-dependent stretching of ZO proteins. ZO-1, ZO-2 are shown schematically, with their structural domains (see Fig. 1). ZO-1 is shown either in a stretched (a, b, c) or folded (d) conformation. DbpA is shown in green (a-c) or yellow (d). Actomyosin filaments are schematically shown as red lines, the number of lines proportional to tension/contractility. Claudin, JAM and occludin proteins are also schematically shown as transmembrane proteins, with their cytoplasmic C-terminal regions interacting with the PDZ1, PDZ3 and GUK regions of ZO-1, respectively. **(a)** Actomyosin tension and heterodimerization maintain ZO proteins in a stretched conformation, promoting DbpA and occludin binding to the ZPSG domain. **(b)** Under lower tension conditions ZO proteins are maintained in the stretched conformation through heterodimerization. **(c)** Upon depletion of ZO-2, ZO-1 is maintained in the stretched conformation when actomyosin tension is higher (e.g. growth of cells in 2D on glass coverslips). **(d)** Lower actomyosin tension (e.g. 3D culture in Matrigel, or treatment of 2D cultures with blebbistatin) combined with expression of only one ZO protein leads to reduced junctional accumulation of occludin and DbpA, and degradation of DbpA. In the presence of the proteasome inhibitor MG132, DbpA is not degraded, but is translocated to the nucleus, where it regulates target gene transcription (Fig. 4a-b).

We propose that the stretched conformation of ZO proteins is the active, conformation, which promotes interaction with DbpA, occludin, and other ligands, through their ZPSG domain. The ZPSG region of ZO-1 has several additional interactors: α-catenin ^16,55^, JAM ^21^, G_α12_ ^56^, Apg-2 ^57^, and Shroom2 ^58^. Future studies should address whether these interactions, and their downstream signaling, can be modulated negatively by ZO-1 folding, and positively by tension-and heterodimerization-induced stretching. According to our model, the folded conformation of ZO-1 and ZO-2 represents their inactive form, in which the ZPSG domain is unable to bind to physiological interactors, due to the competition by the C-terminal domain. Interestingly, some of the ZO-1 interactors, such as α-catenin, are not localized at TJ, but at AJ. Since ZO-1 has been localized at AJ both in cell culture models ^59^ and in tissues ^60^, this raises the possibility that force-dependent modulation of ZO-1 may be important in AJ assembly, and its linkage to the actin cytoskeleton.

FRAP studies show that the association of ZO-1 with the TJ membrane is dynamic, with a soluble cytoplasmic pool in equilibrium with a membrane-associated pool ^61^. It remains to be determined whether the cytoplasmic pool of ZO-1, exchanging with the TJ membrane, as well as monomeric ZO-1 that traffics to the membrane during junction biogenesis, is or is not in a monomeric folded conformation. Our SIM and PLA results indicate that folded ZO-1 can be detected both at junctions and in the cytoplasm, suggesting that folded ZO-1 molecules can remain anchored to the membrane, presumably through interactions of their PDZ1 and PDZ3 domains with claudins and JAM-A, respectively (Fig. 7d). Activation/stretching of ZO proteins, either through heterodimerization and/or through increases in actomyosin contractility (in the physiological range, <5 pN), could be a useful mechanism to coordinate the assembly of the TJ scaffold ^25^ with its linkage to the actin cytoskeleton. Post-translational modifications may also contribute to regulating ZO protein stretching/folding, for example by modulating the affinity of interaction between the C-terminal region and/or the ZPSG with different interactors.

ZO protein stretching and folding in response to extrinsic or intrisic mechanical cues may be relevant to the role of ZO proteins in development and disease. For example, down-regulation of one or both ZO proteins has frequently been observed in cancer (reviewed in ^62^), and substrate stiffness/softness may differentially affect the proliferation and survival of cells expressing one or both ZO proteins. In addition, although epithelial tissues typically express either two or three ZO proteins ^63^, ZO-1 is the only ZO protein expressed in cells of mesodermal origin ^64^ and in extraembryonic mesoderm of mouse embryos ^65^. Thus, the early embryonic lethal phenotype of ZO-1-KO mice ^65^ could be explained by lack of tension-dependent signaling by ZO-1, leading to impaired proliferation of extraembryonic tissues. The early embryonic lethal phenotype of ZO-2-KO mice ^66^ might instead indicate a requirement for ZO-1/ZO-2 heterodimerization in the morphogenesis of a soft embryonic tissue. Conversely, since epithelial cells lacking ZO-2 can populate epithelial organs in chimeric mice ^66^, either ZO-1/ZO-3 heterodimerization ^28^ or higher tension might compensate for the lack of ZO-2 in adult tissues.

In summary, we propose that activation of ZO proteins by stretching is a new mechanism of cross-talk between the contractile cytoskeleton and junctions, through which tensile forces are transduced into downstream signaling by regulating the junctional localization of interactors of the ZPSG regions of ZO proteins. This could be one among different mechanisms through which mechanical forces regulate morphogenesis and homeostasis of epithelial and endothelial tissues.

## ACKNOWLEDGEMENTS

We are grateful to all colleagues who provided cell lines and reagents, and especially to Prof. S. Tsukita for the gift of ZO-1 KO cells. This work was supported by the Canton and Republic of Geneva and the Swiss National Foundation (grant n.31003A_152899 to SC) and the National Research Foundation (NRF), Prime Minister’s Office, Singapore under its NRF Investigatorship Programme (NRF Investigatorship Award No. NRF-NRFI2016-03 to JY).

## AUTHOR CONTRIBUTIONS

D.S. and S.C. conceived and designed the project, analyzed the data, and wrote the manuscript. D.S. prepared constructs and carried out most experiments. S.L. and J.Y. carried out magnetic tweezers experiments. I.M. and L.J. prepared constructs. T.L and D.S performed SIM imaging.

